# AF2BIND: Predicting ligand-binding sites using the pair representation of AlphaFold2

**DOI:** 10.1101/2023.10.15.562410

**Authors:** Artem Gazizov, Anna Lian, Casper Goverde, Sergey Ovchinnikov, Nicholas F. Polizzi

## Abstract

Predicting ligand-binding sites, particularly in the absence of previously resolved homologous structures, presents a significant challenge in structural biology. Here, we leverage the internal pairwise representation of AlphaFold2 (AF2) to train a model, AF2BIND, to accurately predict small-molecule-binding residues given only a target protein. AF2BIND uses 20 “bait” amino acids to optimally extract the binding signal in the absence of a small-molecule ligand. We find that the AF2 pair representation outperforms other neural-network representations for binding-site prediction. Moreover, unique combinations of the 20 bait amino acids are correlated with chemical properties of the ligand.

## 1 Introduction

Accurate prediction of ligandable sites in proteins remains an outstanding challenge. Though some methods take advantage of homology to transfer binding-site annotation between proteins based on structural similarity, [15] [37] true *de novo* binding-site prediction remains difficult. Such a capability could generate novel functional hypotheses across whole proteomes or focus drug-discovery efforts.

In particular, the task of predicting small-molecule binding sites remains challenging. Several machine-learning approaches have been proposed to tackle the problem of small-molecule binding-site prediction [3] [32] [30] [7]. These approaches typically build residue-or atomic-level graph representations of a protein structure [32] [30] or use embeddings from models trained to predict masked sequence tokens [24] or the sequence of a protein given its structure [16] [11] [17]. Small-molecule binding sites are often found in “frustrated” regions of a protein [13], so models that predict sequence from structure might do well to recognize such sites. Models such as ESM-IF or proteinMPNN [16] [11] might detect solvent-exposed hydrophobic clusters of residues or completely buried collections of polar residues for a binding-site prediction. We reasoned that a structure-prediction model, such as AlphaFold2 (AF2) [19], might similarly detect such properties while also providing an additional channel to identify binding sites through pairwise attention between frustrated regions of a target protein and additional amino acids provided for “co-folding”. If we could extract this additional attention signal, AF2 might then offer a rich embedding for the binding-site prediction task.

Because protein intramolecular contacts can closely approximate protein-ligand contacts [28] [22], we reasoned that individual “bait” amino acids could be supplied to AF2, in conjunction with the template of a protein target (either from the protein data bank (PDB) or predicted), to tease out the small-molecule-binding signal. The “bait” might allow AF2 to alleviate regions of frustration in the target template (“finish folding”), and binding-site prediction would then proceed by tracking what AF2 does with this bait. Here, we show that AF2’s internal representation of protein structure and sequence is sufficiently expressive to train a logistic regression model, AF2BIND (AF2 Bait-Informed Neural Descriptor), to accurately predict small-molecule binding-site residues in proteins, without multiple sequence alignments, homology models, or knowledge of the true ligand. Finally, we use AF2BIND to compute ligand polarity compatible with a binding pocket.

### 1.1 AF2BIND: a logistic regression model using AF2 features

AF2BIND is a logistic regression model trained from AF2 pair features to predict each residue’s probability of contacting a small-molecule ligand, given a target protein structure (Fig. 1). The goal in our development of AF2BIND was to understand if features derived from AF2 provide a richer description for the small-molecule binding-site prediction task than previously used models. AF2 has already been used in predicting protein-protein and protein-peptide structures [34] [21] [4], but it is less clear whether it could be useful for predicting small-molecule binding sites because small molecules are not explicitly modeled or accounted for in the training objective. Still, AF2 was trained on proteins that contain small molecules, and it sometimes even predicts the correct rotamers for binding, e.g. in metal or heme sites, so some nuanced binding signatures might be embedded in its features that could be used to train a model for binding-site classification. Given that AF2, trained only on monomeric protein structures, can accurately predict protein-peptide co-structure with no co-evolutionary information of the peptide (and no examples in the training set), we reasoned AF2 might also be useful for predicting small-molecule-binding sites, which often locally resemble amino-acid sidechains [28].

**Figure 1.**
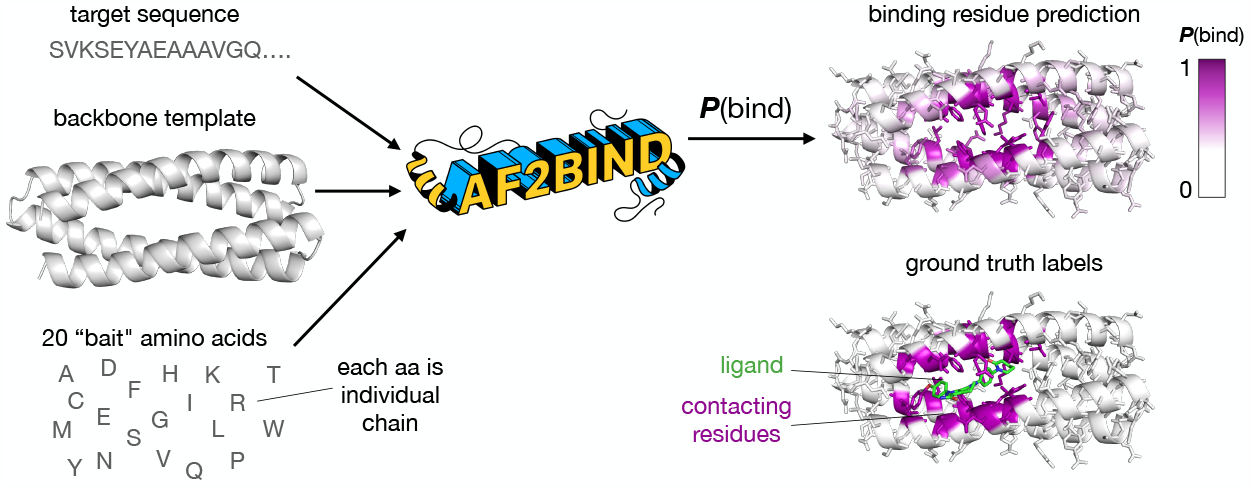
AF2BIND uses features from AlphaFold2 to predict ligand-binding residues in a target protein. The inputs to AF2BIND are the target sequence, target backbone, and 20 bait amino acids, which are surrogates for a small-molecule ligand. The output is a prediction for each residue of the target protein, **P**(bind), which is the probability that the residue is a small-molecule-binding residue. The model is trained on ground-truth labels from a couple thousand non-redundant protein-ligand co-crystal structures from the PDB.

We set out to train a binding-site prediction model that was maximally interpretable, with few operations that transform the AF2 embeddings. We chose a logistic regression model trained directly on AF2’s pairwise representation, which assigns each pair of amino acids in an input sequence a tensor that is used to predict the relative position of these residues in the structure (Fig. 2A). We give AF2 the sequence of the target protein and its backbone structure as a template (We do not supply AF2 with a multiple sequence alignment.). To allow AF2 to “finish folding” the template protein, we also provide twenty disconnected “bait” amino acids as individual chains, by appending these to the target sequence with large residue offsets between each. Each of the canonical twenty amino acids is used. Instead of using multiple recycles, as is common for structure prediction, we only compute a single pass through the AF2 model to generate the pair representation. Our motivation for this is to extract the initial attention between target and “bait” sequence, unbiased by arbitrary placement of “bait” amino acids by the structure module. The pairwise embeddings between each of the twenty bait amino acids and a target residue are concatenated and fed into the logistic regression model for training, with the objective to predict the label for that residue (1 or 0, whether it is a binding residue or not, Fig. 2A). AF2BIND thus takes a target sequence and backbone structure and computes a probability, **P**(bind), for each residue to participate directly in a small-molecule binding site (Fig 1 and see Methods). In addition to the per-residue binding-site prediction, the interpretable nature of the logistic regression model allows us to identify which bait amino acids are contributing to the sigmoidal activation of **P**(bind) (Fig. 2B).

**Figure 2.**
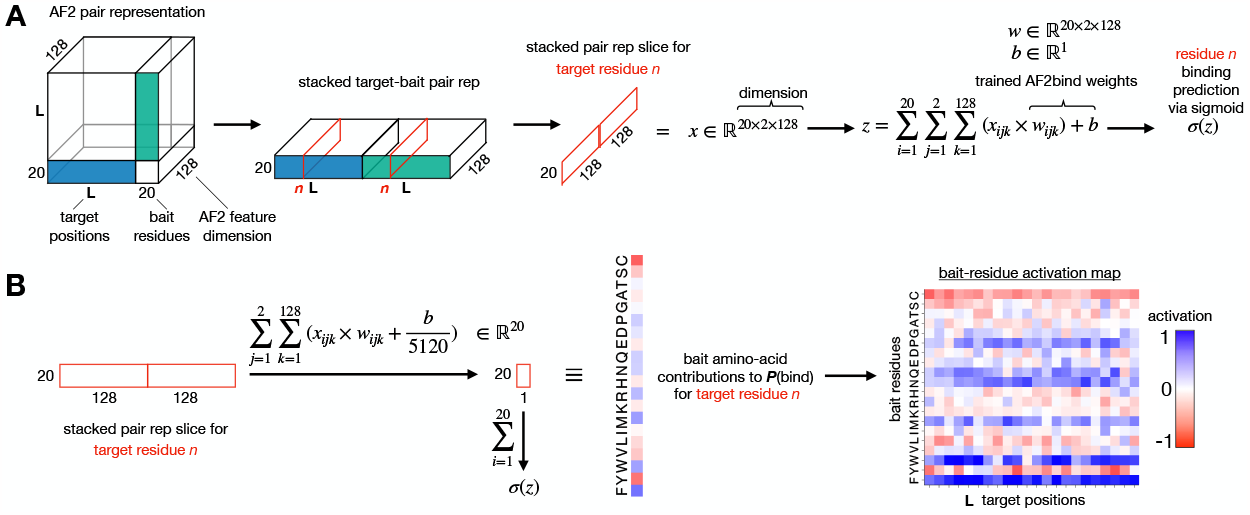
The pair representation of AlphaFold2 is used as input to a logistic regression model, AF2BIND, for prediction of a ligand-binding residue. A) The blue and green tensors represent the pairwise attention between target and 20 bait amino acids; these features are extracted, flattened and fed into a logistic regression model for binding-site prediction, **P**(bind). B) Since the features directly map to the 20 bait amino acids, the contribution (activation) of each amino acid to the binding-residue prediction can be extracted.

## 2 Results

### 2.1 AF2 provides superior embeddings for binding-site prediction

Direct comparison of AF2BIND to other ligand-binding-site predictors is challenging because most published train/test data-splits contain significant amounts of data leakage. Instead, we compare the performance of different representations from various pre-trained models for the binding-site prediction task on our train/test split (see Dataset in Methods). We compared the performance of AF2BIND with the single representation features from AF2, ESM2 [24] and ESM1-IF [16] for just the target sequence or structure (no bait amino acids were included). Recently, ESM2 [2] and ESM1-IF [3] representations were used for binding-site prediction on a different train/test set. We find the pair representation of AF2 outperforms these other representations for the small-molecule binding-site prediction task (Fig. 3).

**Figure 3.**
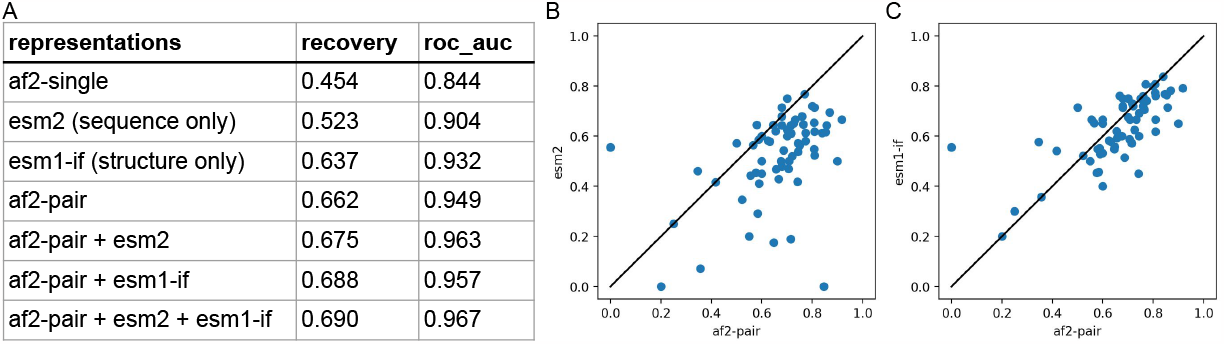
The pair representation of AF2 is most effective at binding-residue prediction. A) The average binding-site recovery on (10-fold cross) validation set given different representations as input to logistic regression. B) Binding-site recovery of test set between B) ESM2 and AF2-pair representations and C) ESM1-IF and AF2-pair representations. Each point is a different target protein.

To assess model performance, we compute the fraction of binding-site residues recovered from the top predictions, sorted by highest to lowest **P**(bind) (See Methods for details). While the pair representation of AF2 provides the best singular embeddings for the binding-site prediction task (Fig. 3), a combination of AF2-pair and ESM embeddings provides the best recovery and AUC ROC metrics (Fig. 3A). The average recovery for binding-residue prediction was 66% for AF2-pair representation (AF2BIND) and increased only slightly to 69% when combined with ESM2 and ESM1-IF.

### 2.2 Exemplar performance of AF2BIND on held-out protein classes

AF2BIND can accurately predict binding residues of rigorously held out protein classes. We held out G-protein coupled receptors and bromodomains (as well as other classes) from training and validation, based on sequence similarity and structural similarity (see Methods). AF2BIND confidently predicts the binding residues of these proteins (Fig. 4). Not every residue in the binding site is predicted with equal probability, providing a hierarchy of residues most likely involved in binding a small molecule. This ranking scheme gives additional information over volumetric pocket-finding algorithms [23], as it suggests which residues might preferentially engage a ligand. **P**(bind) is not trivially correlated with conservation (Fig. S2), as shown by the low conservation in the orthosteric binding site of the mu opioid receptor relative to the more conserved intracellular core and G-protein-binding regions (Fig. S2A). Furthermore, slight variations in the protein backbone lead to reproducible predictions (Fig. S2B). For example, four structures of the human mu opioid receptor were compared, with average backbone root mean square deviation (RMSD) of 0.7Å. The average spread (standard deviation) in **P**(bind) for each residue was very low (0.02), and the variance did not correlate with the magnitude of **P**(bind). Because we mask the sidechain dihedral angles of the input template of the target protein, AF2BIND is insensitive to the sidechain rotamers of the target protein, which is advantageous if the rotamers are uncertain (common for predicted protein structures). Indeed, we found that masking the sidechain dihedrals of the holo structures during training led to equal (or even slightly better) predictive power (recovery) of the model, relative to a model trained with the additional sidechain information given to AF2 (Fig. S1). Because one of the most common differences between unbound and bound structures of proteins is the change of a rotamer in a binding-site residue [8], this insensitivity to sidechain coordinates should prove beneficial when analyzing unbound or ambiguous structures. Moreover, changes in backbone coordinates between unbound and bound states are on average small (RMSD < 1Å), within the range where predictions by AF2BIND are robust.

**Figure 4.**
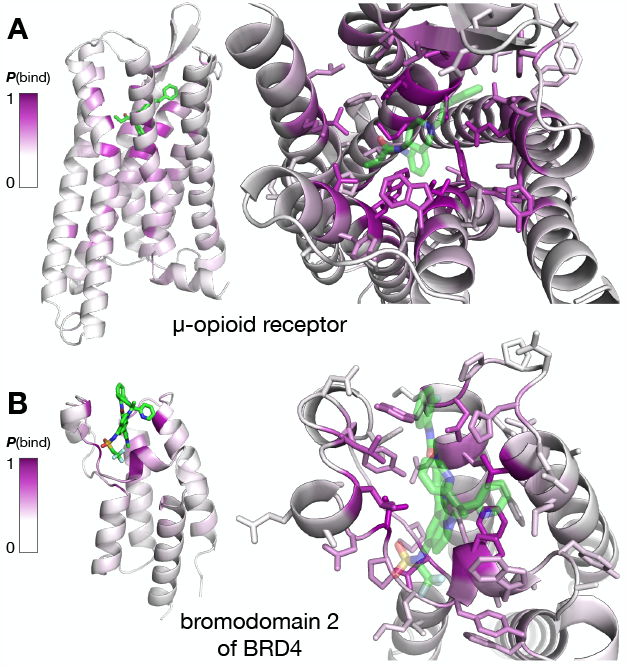
AF2BIND makes accurate predictions on held-out proteins. A) Per-residue binding-site predictions of the human mu opioid receptor (pdb 8ef5) bound to fentanyl and of B) the second bromodomain of human BRD4 (pdb 7ruh) bound to an inhibitor. AF2BIND never saw a G-protein coupled receptor, bromodomain, or similar proteins during training. The highly accurate predictions were made without knowledge of the bound ligand. The color scale of the binding probability, P(bind), scales from white (P(bind) = 0) to purple (P(bind) = 1).

### 2.3 Contributions of bait residues correlate with ligand hydrophobicity

Because AF2BIND uses a linear combination of AF2 pair features, the contribution of each of the twenty bait amino acids to the binding-residue prediction can be uniquely attributed (Fig 2B). The ease of this assignment is a distinct advantage of the logistic regression model. We analyzed the propensity of the “bait” amino acids to contribute to **P**(bind) as a function of polarity (fraction of carbon atoms) of the true ligand (Fig 5). We found that certain bait-residue combinations were negatively and positively correlated with ligand hydrophobicity (Fig 5, A and B, respectively). This activation analysis represents a possible ligand as a linear combination of the twenty amino acids. Hydrophobic bait amino acids predominately contribute to the binding-site prediction of the supernatant protein factor, 4omj, which binds the hydrophobic terpenoid, 2,3-oxidosqualene (Fig 5C). On the contrary, hydrophilic bait amino acids are the main contributors to binding-site prediction for the 4-diphosphocytidyl-2C-methyl-D-erythritol kinase, 2v2z, which binds the polar substrate, 4-diphosphocytidyl-2C-methyl-D-erythritol (Fig. 5D).

**Figure 5.**
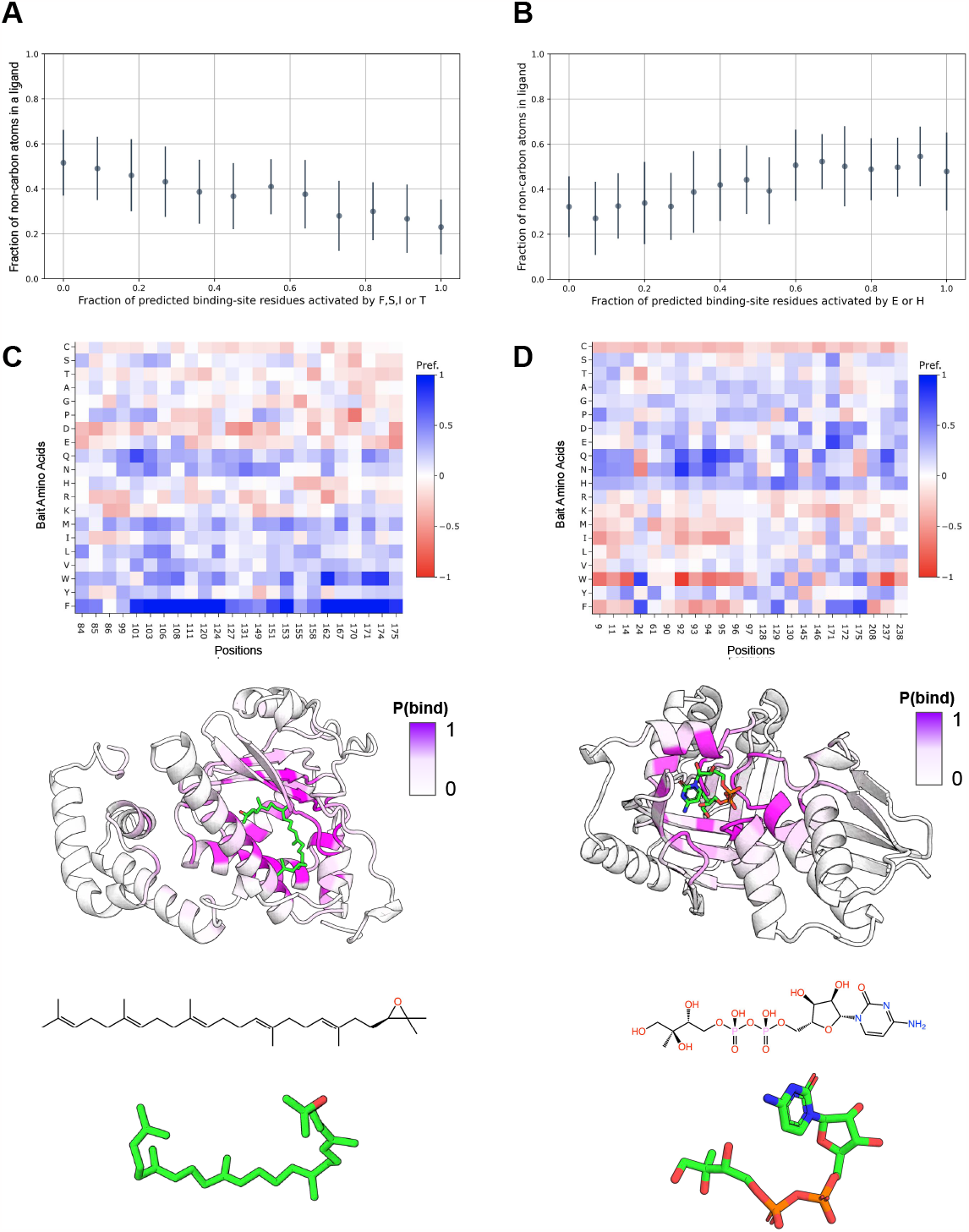
Activations of the bait amino acids (and their combinations) are correlated with ligand hydrophobicity. The fraction of hydrophobic amino acids FSIT activated (A) and the fraction of hydrophilic HE (B) had the highest absolute pearson correlation relative to the fraction of non-carbon atoms in the respective ligands. C) Analysis of predictions for transport protein (4omj). The ligand (green) has a high fraction of carbon atoms, and the dominant amino acids that were activated among the bait residues are W and F. On the other hand when the fraction of carbon atoms in a ligand decreases (2v2z), the activation value of W and F drops, and that of polar amino acids (such as Q and N) increases.

## 3 Discussion

Here, we showed that a deep neural network, AF2, trained on the task of single-chain protein-structure prediction, has the surprising ability to discern small-molecule-binding sites, without needing to provide a sterically or chemically compatible ligand. The pair representation of AF2 provides an effective embedding for the training of a dedicated logistic regression classifier, AF2BIND, that can accurately assign small-molecule-binding labels to each residue in a target protein (66% recovery). We used twenty individual-chain amino-acid “bait” residues to effectively tease out a ligand-binding signal from the pair representation of AF2. An activation analysis of the bait residues, which are built from each type of amino acid, correlates with ligand hydrophobicity.

To train AF2BIND, we created a rigorously split dataset of protein-ligand complexes curated from the entire PDB. The split was based not only on sequence but also on fold, evolutionary classification, and binding pocket similarity. This rigorous split is important for the binding prediction task because binding sites are highly conserved in proteins, even if the overall sequence diverges. Our motivation was to prevent any leakage of the training data into the test data so we could accurately determine the ability of the model to generalize for *de novo* binding-site prediction. After creating this split, we found that regularization was needed to keep from overfitting the model. The regularized model should therefore be well-calibrated, meaning the recovery is correlated with the magnitude of **P**(bind). In initial stages of training the model (before we filtered for data quality such as real-space correlation coefficient of the ligand), we looked at particular data points that were consistently predicted with low recovery and found that the quality of the crystallographic data for these binding sites was poor. Thus, AF2BIND might also be used to determine if a ligand placement might be spuriously modeled into experimentally determined structures.

Because our main motivation was probing the inherent binding-site prediction capabilities of AF2, our logistic regression model is purposefully simple to maximize interpretability; and there likely remains some room for model improvement. The theoretical ceiling for recovery in this prediction task is unclear because the true ligand is not used for the prediction, so the ceiling is likely well below 100% recovery. Simply clustering the predictions by proximity to neighboring residues with high **P**(bind) could increase performance. Lastly, the use of a limited number of bait amino acids by AF2BIND could result in a dilution of the binding signal as the number of binding sites in a target protein grows (AF2BIND was trained on single-chain proteins up to 500 amino acids.). Offsetting any signal dilution through, e.g., increasing the number of bait amino acids is one possibility. Altogether, there are many rich avenues for future exploration.

The ability to accurately predict *de novo* binding sites could enable binding-site identification across large swaths of newly predicted protein structures. Homology modeling approaches such as AlphaFill rely on “filling” predicted proteins with ligands taken from crystal structures of homologous proteins.[25] [15] While useful, these approaches are not applicable to new folds or unliganded sites in the PDB. Other prediction approaches either rely on achieving the correct chemical compatibility of the ligand to identify a site (blind docking) [5] [9] [14] or are trained to predict voxels that carve out the space occupied by a known ligand. [12] [18] The myrid unliganded structures now available in AlphaFoldDB [36] and ESM Metagenomic Atlas [24], combined with the predictive power of AF2BIND, offer tantalizing opportunities to discover novel binding sites across the tree of life.

## 4 Methods

### 4.1 Extracting AF2 features for binding-residue prediction

AF2 is trained to output a protein structure given an input protein sequence. Along with the sequence, AF2 can take as optional input a multiple sequence alignment (MSA) and structural template. These inputs get transformed internally by the model through several attention layers to a “single” and a “pair” representation, so-called because there is an embedding for each single residue or each pair of residues of the input sequence, respectively. The residue identity of each amino acid, as well as its relative position within the input sequence, affect its precise numerical embedding by AF2, which uses the embeddings within the structure module to compute the 3D coordinates of the protein. We tested the usefulness of the single and pair representation of AF2 for binding-residue prediction. To bias the single and pair features to the task of ligand binding, we append 20 disjoint (by a large residue-index offset) amino acids to the sequence of a target protein. These 20 amino acids (one for each of the canonical residue types) act as “bait” ligands for AF2. We use as a template the structure of the target protein (no template was provided for the 20 “bait” amino acids.) and found that ablating the sidechain dihedral information beyond C*β* atom was most effective for downstream binding predictions. Because we use a template of the target protein, we do not need to use a MSA, as AF2 will output a structure that is practically identical to the input template if the sequence of the template and the input sequence are the same [29]. AF2BIND uses the pair representation between a residue *j* in the target protein and each of the 20 bait amino-acids (Fig. 2A). The concatenated pair representations (dimensionality of 5120 or 20x2x128 features) are input to AF2BIND for training the logistic regression model. AF2BIND is trained to predict that a given residue in the target protein is involved in binding a ligand; we call the sigmoidal activation **P**(bind) for probability of binding.

### 4.2 Dataset

We trained AF2BIND on single-chain protein-ligand complexes filtered from all crystal structures in the protein data bank (PDB) accessed in March 2023. The PDB was first filtered by resolution (< 3.6Å), R factor (< 0.35), chain length (between 40 and 500 residues), monomeric oligomerization state, and absence of RNA/DNA polymers. The single-chain proteins needed to have a small-molecule ligand bound with buried surface area > 100Å^2^. The ligand needed to contain between 10 and 200 heavy atoms, and peptides were excluded. Proteins contacting a ligand with more than one chain in the asymmetric unit or crystal lattice were excluded. We removed ligands that were covalently bound (except for porphyrins) or listed as common crystallization additives. For obtaining binding-residue labels to train AF2BIND, we consider only ligands with real-space correlation coefficients > 0.85, real space R value < 0.25, and average occupancy > 0.9. Approximately 14k PDB files (15k chains) remained, containing roughly 18k ligands. We clustered these using mmseqs2 [33] by 30 percent sequence similarity and 80 percent coverage (1280 clusters). We also clustered the proteins structurally using foldseek [35] and combined the foldseek and mmseqs2 clusters. To create our final dataset for training, we extracted up to 3 proteins from each cluster, ranked by number of residues in the binding pocket (max), diffraction resolution (min), and R-free of the structure (min). Within a cluster, preference was also given to proteins with different ligand binding-site locations. The resulting dataset consisted of 1902 proteins and 2110 ligands. Binding residues were determined based on a maximum heavy-atom distance of 5Åbetween ligand and residue. We subsequently augmented binding-residue labels of these proteins using the 15k chains if a chain had TM-score > 0.8 and > 90 percent sequence similarity in the binding residues.

### 4.3 Training

AF2BIND is a logistic regression model trained for 320 epochs and a batch size of 12 proteins, using the ADAM optimizer with learning rate of 1e-4. The input features were normalized by the mean and standard-deviation of the training set. To down-weight redundancy, each sample was compared to all other samples in the training set, and was weighted by 1 / number of proteins > 0.5 TM-score. To prevent over-fitting, we scanned L2 regularization weights and selected a weight (0.03) where the average train and validation recovery were about the same (Fig S3). This was performed across 10 cross-validation splits. We tested the performance of the final weights on a test set of about 70 protein–small-molecule complexes (Fig 3B, C).

### 4.4 Representations

For all representations evaluated, we used the default representations provided by ESM2-3B, ESM1-IF and AlphaFold2, which are extracted from the final layer before the prediction task. When combining representations, as in (Fig 3A), we concatenate the representations and train a new model.

### 4.5 Assessment Metric

For comparing performance of different representations, we adopt a metric often used for assessing contact accuracy in proteins [20]. In short, the predictions are sorted by most to least confident, and the fraction of the top *N* predictions are evaluated, where *N* is the number of expected correct labels. The advantage this approach is that it allows for comparisons of methods where the confidence metrics maybe in different ranges or not fully calibrated. We adopt this metric for our binding site recovery evaluation in (Fig 3A).

### 4.6 Robust splitting of training/validation and test

Since our goal was to develop a method that can generalize to unseen folds, we took special care to make sure there was no structural overlap between training, validation, and test set. This was done by creating 11 independent sets of about 20 proteins each. We computed all-by-all TM-scores for the 2k proteins in the dataset. The proteins were sorted from least similar to most similar by TM-score [38] to all other proteins in the set. The proteins were then assigned one by one to each of the 11 sets. The 11th set was further modified to include the follow PDB chains: 4lbs_A, 7xn9_A, 4n03_A, 3hlg_A, 7ruh_A, we wanted to exclude these from training for detailed analysis later. At each step, we verified there was no overlap to any of the other proteins already assigned in each set. Proteins were deemed overlapping if their TM-score > 0.5, or they shared any ECOD (topology level) [6], CATH (at topology) [31], SCOP2B (superfamilies level) [1], PFAM [26] or InterPro annotation [27]. The latter three were extracted from the SIFTS database [10]. We also extracted binding-site residues from each protein and used TM-align to compute an all-by-all TM-score for each pair of pockets in the dataset. We used the harmonic mean of the two TM-scores computed for each pair and excluded proteins from test that had similar pockets (harmonic mean of TM-score > 0.6) to any in train. After initial split, the last bin was reserved as the test set. The remaining 10 sets were used to create 10 cross-validation sets, where 9 sets are used for training and 1 set is used for validation. For each cross-validation assignment, proteins that were excluded from the initial split were added back into each train/validation/test set if they were non-redundant and shared similarity within the set but not between sets. A protein was deemed redundant if TM-score > 0.8 and more than 60% of binding sites overlapped. After enrichment, each cross-validation set had on average 600 proteins for train, 30 for validation, and 70 for test. The full set can be downloaded at: https://github.com/sokrypton/af2bind

## 5 Code Availability

Google Colab notebook to run AF2BIND for binding-site prediction and activation analysis can be found here: https://colab.research.google.com/github/sokrypton/af2bind/blob/main/af2bind.ipynb

## A Supplementary Figures

**Figure S1:**
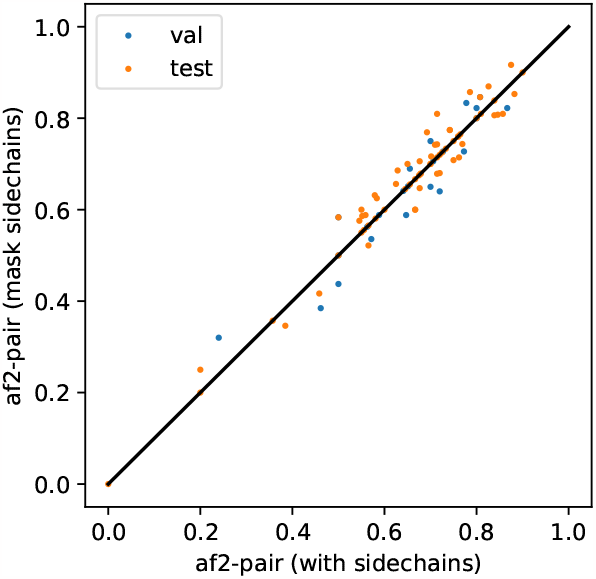
Masking sidechains from template input features does not significantly affect the quality of AF2-pair representations for the task of ligand binding-site prediction. Each dot is a separate protein from either validation (blue) or test (orange) set. X and Y-axis are the binding site recovery values for each protein.

**Figure S2:**
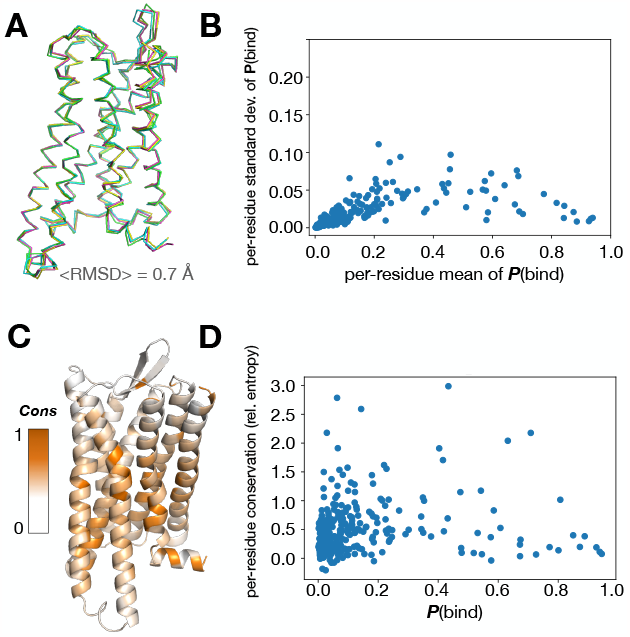
Small changes in protein backbone do not significantly affect **P**(bind). A) Several structures of the human mu opioid receptor (pdbs 8ef5_R, 8ef5_M, 8efq_R, and 8efb_R), with average root mean square deviation of C-alpha atoms of 0.7Å. Predictions by AF2BIND on each of these ensemble members were averaged at the residue level. The plot displays the standard deviation and mean of these predictions. **P**(bind) changes by at most 0.1 but on average by only 0.02, the magnitude of which is uncorrelated with the mean value of **P**(bind). B) **P**(bind) is uncorrelated with amino-acid conservation, as measured by the relative entropy computed from a multiple sequence alignment, using the amino-acid background frequency in the PDB as the reference distribution.

**Figure S3:**
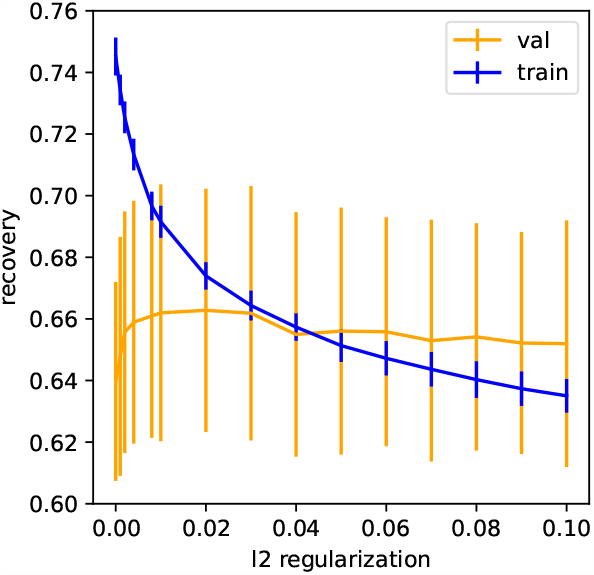
Recovery of binding site on training set and validation set, given L2 regularization, using AlphaFold2 pair representation.

